# PARylation regulates stress granule dynamics, phase separation, and neurotoxicity of disease-related RNA-binding proteins

**DOI:** 10.1101/396465

**Authors:** Yongjia Duan, Aiying Du, Jinge Gu, Gang Duan, Chen Wang, Zhiwei Ma, Le Sun, Beituo Qian, Xue Deng, Kai Zhang, Kuili Tian, Yaoyang Zhang, Hong Jiang, Cong Liu, Yanshan Fang

## Abstract

Mutations in RNA-binding proteins localized in ribonucleoprotein (RNP) granules, such as hnRNP A1 and TDP-43, promote aberrant protein aggregations, which are pathological hallmarks in neurodegenerative diseases including amyotrophic lateral sclerosis (ALS) and frontotemporal dementia (FTD). Protein posttranslational modifications (PTMs) are known to regulate RNP granules. In this study, we investigate the function of PARylation, an important PTM involved in DNA damage repair and cell death, in RNP-related neurodegeneration. We reveal that PARylation levels are a major regulator of the dynamic assembly-disassembly of RNP granules, and the disease-related RNPs such as hnRNP A1 and TDP-43 can both be PARylated and bind to PARylated proteins. We further identify the PARylation site of hnRNP A1 at K298, which controls the cytoplasmic translocation of hnRNP A1 in response to stress, as well as the PAR-binding motif (PBM) of hnRNP A1, which is required for the delivery and association of hnRNP A1 to stress granules. Moreover, we show that PAR not only dramatically enhances the liquid-liquid phase separation of hnRNP A1, but also promotes the co-phase separation of hnRNP A1 and TDP-43 *in vitro* and their interaction *in vivo.* Finally, we establish that both genetic and pharmacological inhibition of PARP mitigates hnRNP A1 and TDP-43-mediated neurotoxicity in cell and *Drosophila* models of ALS. Together, our findings indicate a novel and crucial role of PARylation in regulating the assembly and the dynamics of RNP granules, and dysregulation of PARylation may contribute to ALS disease pathogenesis.

## INTRODUCTION

Eukaryotic genomes encode a large number of RNA-binding proteins that can associate with RNAs to form ribonucleoprotein (RNP) complexes. RNPs contain conserved RNA binding domain(s) and protein-protein interaction domain(s). They are presented in both nucleus and cytoplasm, where they play a major role in RNA homeostasis including RNA processing, transport and turnover (Anko and Neugebauer, 2012; Lunde et al., 2007; Muller-McNicoll and Neugebauer, 2013). RNPs can form granules by liquid-liquid phase separation (LLPS), and aberrant RNP granules enriched of irreversible amyloid fibrils and aggregations can lead to the pathogenesis of human neurodegeneration diseases including amyotrophic lateral sclerosis (ALS) and frontotemporal dementia (FTD) (Lin et al., 2015). Posttranslational modifications (PTMs) such as phosphorylation, ubiquitination and acetylation are known to regulate the assembly and function of RNP granules (Brady et al., 2011; Cohen et al., 2015; Dammer et al., 2012; Li et al., 2017). Recently, methylation and phosphorylation are reported to modulate phase transition of hnRNPA2 and FUS (Hofweber et al., 2018; Qamar et al., 2018; Ryan et al., 2018; Luo et al., 2018). However, the function of other important PTMs in regulating LLPS and RNP granules remains to be explored.

Poly(ADP-ribosyl)ation (PARylation) is a reversible PTM process by which poly(ADP-ribose) (PAR) polymerases (PARPs) add ADP-ribose (ADPr) units to the Glu, Asp, Lys, Arg or Ser residue of a protein (Martello et al., 2016; Zhang et al., 2013) whereas enzymes such as PAR glycohydrolase (PARG) remove them (Niere et al., 2012; Slade et al., 2011). The opposing effects of PARPs and PARG in regulating protein PARylation play an important role in a variety of cellular functions, including chromatin remodeling, DNA repair, transcription regulation and cell death (Ahel et al., 2008; Andrabi et al., 2006; Frizzell et al., 2009; Singh et al., 2017). Dysregulation in PARylation is therefore involved in various disease conditions such as cancer, neurodegeneration, oxidative stress, neural injury, and regeneration (Deng, 2009; Hanai et al., 2004; Martire et al., 2015; Brochier et al., 2015). Interestingly, PAR as well as some PARPs and PARG are found in cytoplasmic stress granules (SG) and may regulate microRNA-mediated translational repression and mRNA cleavage (Gagné et al., 2008; Kotova et al., 2009; Leung et al., 2011).

Mutations in several RNA-binding proteins such as heterogeneous nuclear ribonucleoprotein A1 (hnRNP A1, encoded by *HNRNPA1*) and TAR DNA binding protein 43 kDa (TDP-43, encoded by *TARDBP*) are known to promote protein aggregation and cause familial ALS (Kim et al., 2013; Neumann et al., 2006). In this study, we show that PARylation plays an important role in regulating SG dynamics, phase separation, protein-protein interaction, and the neurotoxicity of hnRNP A1 and TDP-43, and RNAi knockdown of PARP1 or application of a PARP inhibitor significantly suppresses hnRNP A1 and TDP-43-mediated neurodegeneration, implicating potential therapeutic values of PARP inhibitors to treat ALS and other related diseases.

## RESULTS

### The levels of PARylation regulate the dynamics of SGs that contain ALS-related RNPs

Although PARylation occurs primarily on PARP proteins, association of PAR with SGs and ALS-related RNPs was observed (Gagné et al., 2003; Gagné et al., 2008; Leung et al., 2011). To determine whether PARylation levels regulated SGs and whether known disease-related RNPs such as TDP-43 and hnRNP A1 could be affected, we tested how Olaparib, a Food and Drug Administration (FDA)-approved PARP inhibitor targeted for cancer treatment (Fong et al, 2009), impacted on the response of TDP-43 and hnRNP A1 when cells were stressed (arsenite, 100 μM). SGs were examined by immunocytochemistry using the SG marker T-cell-restricted intracellular antigen 1 (TIA-1)-related protein (TIAR) (Gottschald et al., 2010). TDP-43 and hnRNP A1 were recruited to SGs when cells were stressed, and inhibition of PARP by Olaparib significantly delayed the assembly of SGs as well as the translocation of TDP-43 (Fig. 1A-1C) and hnRNP A1 (Fig. S1A-S1C) to SGs. Next, we downregulated the PAR hydrolysis enzyme PARG with small interfering RNA (siRNA) to increase PARylation levels in the cells. SGs were induced by arsenite treatment for 30 min, followed by the washout experiment to determine the kinetics of SG disassembly as stress was released (Fig. 1E). Increase of PARylation levels by si-PARG drastically delayed SG disassembly as well as the retrieval of TDP-43 (Fig. S1A-S1C) and hnRNP A1 (Fig. 1D-1F) from SGs. Thus, PARylation levels are a major factor regulating the assembly-disassembly dynamics of SGs associated with ALS-related RNPs.

**Figure 1.**
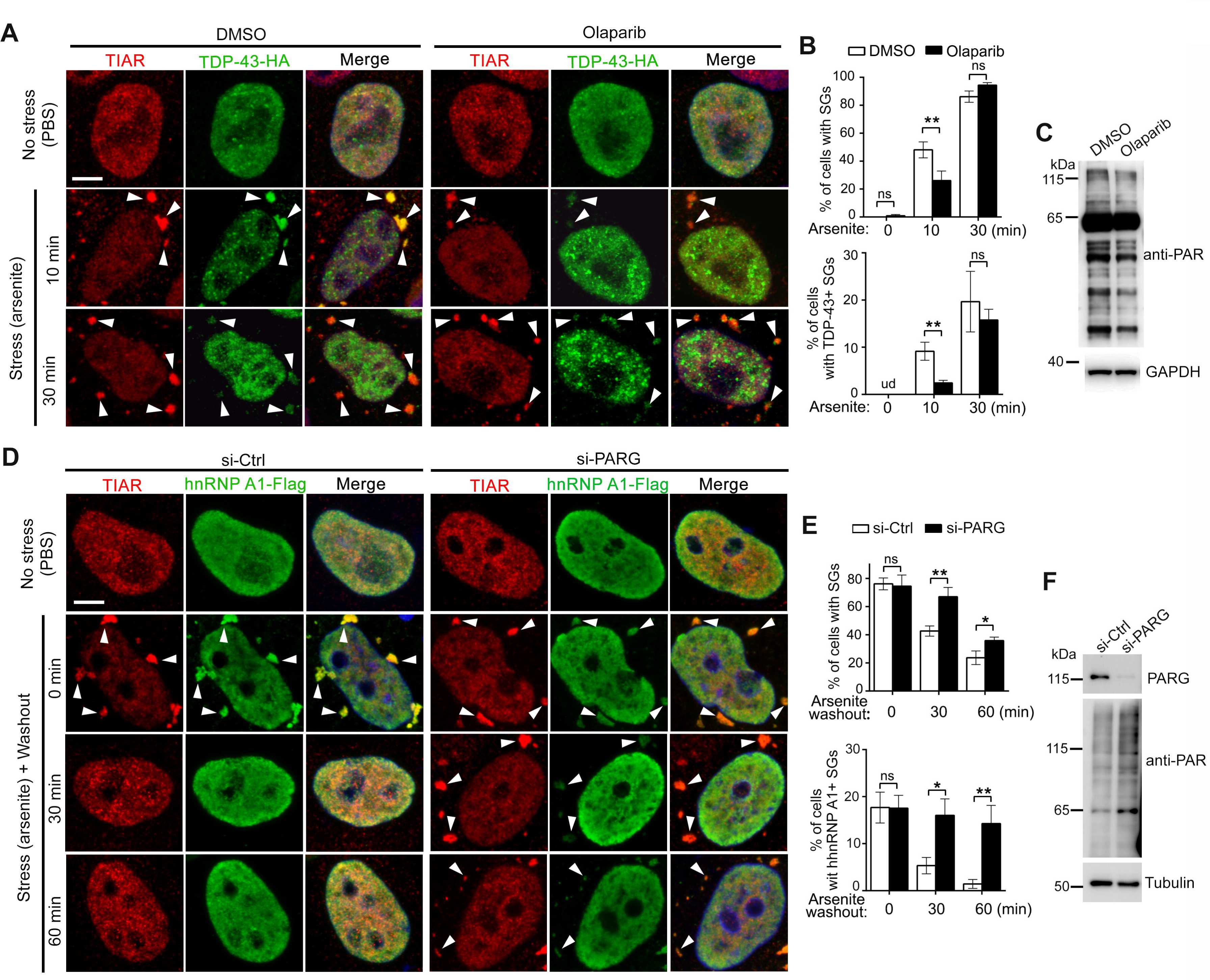
PARylation levels influence the assembly-disassembly dynamics of SGs containing ALS-related RNPs. (**A-C**) PARP inhibition suppresses SG assembly. (**A**) Representative confocal images of HeLa cells treated with PBS (no stress) or 100 μM arsenite (stress) for indicated time in the absence (DMSO) or presence of the PARP inhibitor Olaparib (20 μM). All cells are transfected with TDP-43-HA (anti-HA) and stained for SGs by anti-TIAR (arrowheads), and the merge images with DAPI staining of DNA (blue) are shown. (**B**) The percentage of cells with SGs and TDP-43+ SGs from (A) is quantified. (**C**) Western blot analysis confirms the decrease of overall PARylation levels by Olaparib, with GAPDH as a loading control. (**D-F**) Hydrolysis of PAR by PARG is required for SG disassembly. (**D**) Cells transfected with siRNA (si-PARG) or scrambled siRNA (si-Ctrl) are stressed with arsenite (100 µM) for 30 min and then washout for indicated time. All cells are transfected with hnRNP A1-Flag (anti-Flag) and stained for SGs by anti-TIAR (arrowheads), and the merge images with DAPI staining of DNA (blue) are shown. (**E**) Percentage of cells with SGs or hnRNP A1+ SGs from (D). (**F**) Western blot analysis confirms PARG KD and increase of overall PARylation levels in the cells by si-PARG. Data are shown as mean ± SEM; n = over 100 cells each condition, pooled results of 3 independent repeats; \**p* < 0.05, \*\**p* < 0.01; ns, not significant; ud, undetected; Student’s *t*-test. Scale bars: 10 μm.

### PARylation and PAR-binding of TDP-43 and hnRNP A1

Since PARylation levels affected the dynamics of the RNP granules containing TDP-43 and hnRNP A1, we examined whether these proteins were PARylated or associated with PAR in cells. HA-tagged TDP-43 or Flag-tagged hnRNP A1 was expressed and immunoprecipitated from HeLa cells with anti-HA or anti-Flag, and then examined by Western blotting with a pan-PAR antibody (anti-PAR) that recognizes both mono-ADPr and poly-ADPr. Although there were proteins co-immunoprecipitated with TDP-43 that were immunoblotted positively with anti-PAR, we did not detect obvious PARylation of TDP-43 *per ser* (Fig. 2A). To boost the PARP1 activity in cells, we treated the cells with H_2_O_2_ (500 μM, 10 min) to induce PARP1 activation (Martello et al., 2016), which markedly increased the PARylation levels of hnRNP A1 (Fig. 2B). However, no PARylation of TDP-43 was detected even under this condition. Nevertheless, this result did not exclude the possibility that PARylation of TDP-43 in cells occurred at a very low level that was below our detection sensitivity. In contrast, hnRNP A1 immunoprecipitated from HeLa cells showed robust stead-state and induced levels of PARylation. In addition, there were multiple PAR^+^ bands of proteins of higher molecular weights that were co-immunoprecipitated with hnRNP A1, especially when treated with H_2_O_2_ (Fig. 2B). These were likely PARylated proteins that were associated with hnRNP A1 in the cells.

**Figure 2.**
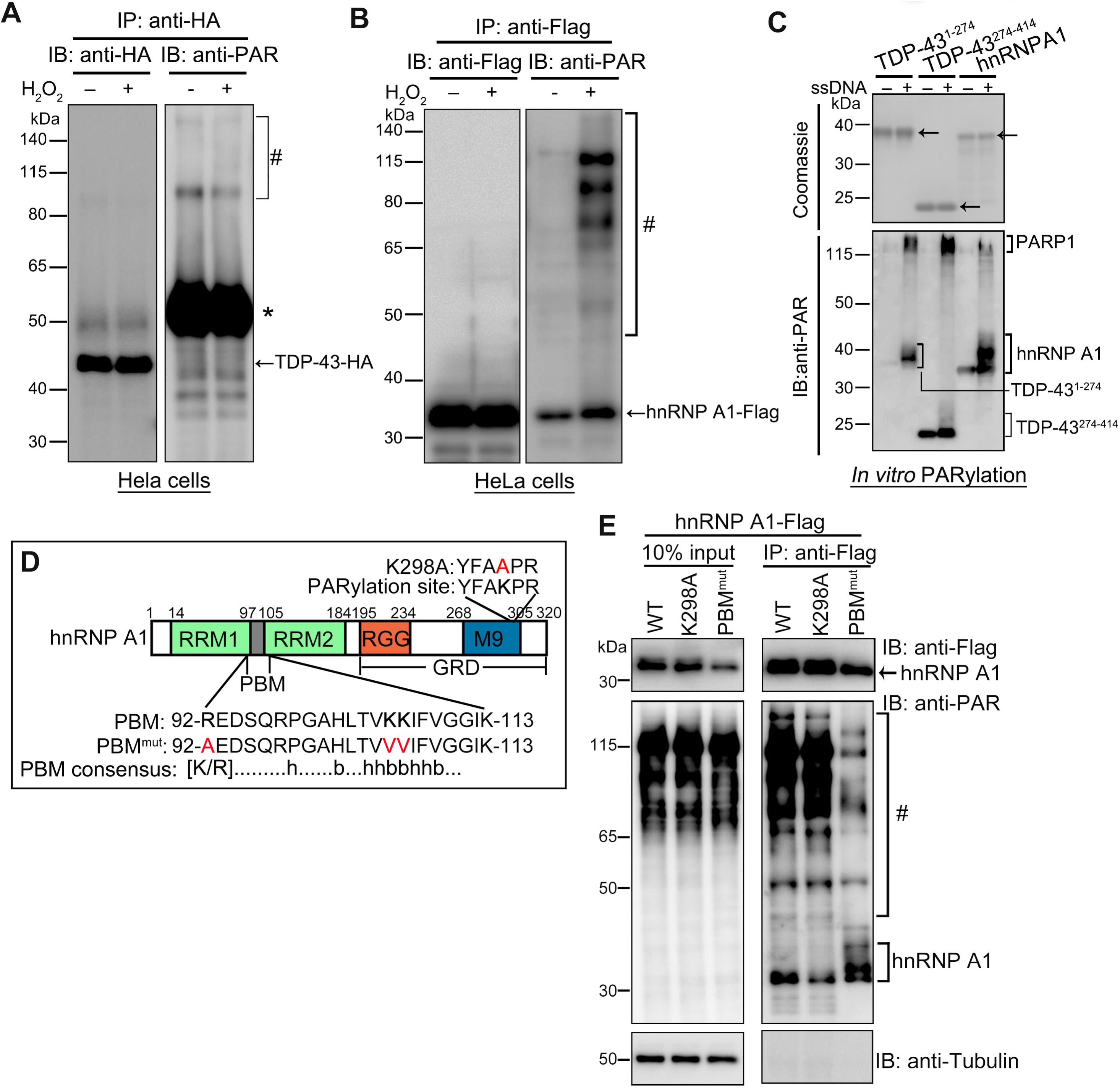
PARylation and PAR binding of TDP-43 and hnRNP A1 *in vitro* and in cells. (**A-B**) HA-tagged TDP-43 (A) or Flag-tagged hnRNP A1 (B) is expressed and immunoprecipitated from HeLa cells that are treated with or without H_2_O_2_, and examined by Western blotting with indicated antibodies. hnRNP A1 shows both steady state and induced levels of PARylation, whereas no PARylation of TDP-43 was detected even when the blot is overexposed to facilitate the detection. (**C**) Recombinant purified His-tagged TDP-43^1-274^, TDP-43^274-414^ and full-length hnRNP A1 proteins (4 μg) were subjected to *in vitro* PARylation assay and then separated on SDS-PAGE gel. Coomassie Brilliant Blue staining confirms equal protein loading (upper panel, arrows indicate the substrate proteins), followed by immunoblotting with anti-PAR (lower panel). ssDNA was added in the *in vitro* reaction system to activate PARP1. The PARylated bands and smears of recombinant TDP-43 and hnRNP A1 as well as PARP1 are indicated. (**D**) Functional domains of human hnRNP A1 protein. RRM: RNA recognition motif; RGG: RGG box RNA binding domain; M9: a nuclear targeting sequence termed as M9; GRD: glycine-rich domain; PBM: PAR-binding motif. The consensus sequence and the residues mutated in the PBM and putative PARylation site are indicated. h, hydrophobic amino acid residues; b, basic amino acid residues (Ji and Tulin, 2009). (**E**) Confirmation of PARylation or PAR-binding deficit of K298A and PBM^mut^ compared to WT hnRNP A1. hnRNP A1-Flag was immunoprecipitated from HeLa cells with anti-Flag and examined by Western blotting with anti-PAR. The blot is overexposed to facilitate the detection of less abundant PARylated proteins. Asterisk, the rabbit anti-HA IgG heavy chain; #, PARylated proteins co-immunoprecipitated with hnRNP A1.

Next, we assessed whether these proteins could be PARylated *in vitro*. Of note, full-length TDP-43 (TDP-43-FL) protein expressed in bacteria was extremely insoluble. We tried several different expression vectors with different purification tags and induction temperatures, but failed to produce sufficient amount of soluble TDP-43-FL protein for the *in vitro* assays (Fig. S3A-S3E). We thus took an alternative approach and divided TDP-43 into two truncations, TDP-43^1-274^ and TDP-43^274-414^ (Fig. S3A and S3F-S3G). The purified His-tagged TDP-43 truncations as well as full-length hnRNP A1 were then subject to the *in vitro* PARylation reaction. Single strand DNA (ssDNA) was added to activate PARP1 in the *in vitro* system and the PARylation levels were examined by SDS-PAGE and immunoblotting with an anti-PAR antibody. ssDNA mimics DNA single-strand breaks that are the most common form of DNA damage in cells and can induce PARP1 activation (Kim et al., 2004; Poirier et al., 1982), which has been frequently used in *in vitro* PARylation assays. Indeed, activation of PARP1 by ssDNA was evident by the dramatic increase of PARylation of PARP1 itself (Fig. 2C). With PARP1 activation, TDP-43^1-274^ and hnRNP A1 showed remarkable PARylation bands and up-shifting smears due to heterogeneity in the length of ADPr polymer attached. The purified TDP-43^274-414^ protein showed a basal level of PARylation (possibly occurred during expression in bacteria), which was slightly increased in the *in vitro* PARlation assay. The PARylation smear of TDP-43^274-414^ was much less intense than TDP-43^1-274^, while hnRNP A1 exhibited the most robust PARylation smear in the *in vitro* assay (Fig. 2C).

The human hnRNP A1 protein contains two closely-related RNA recognition motif (RRMs) in the N-terminal region and a low complexity (LC), glycine-rich domain (GRD) in the C-terminal region that includes an RGG box RNA binding domain and a M9 nuclear targeting sequence (Fig. 2D; He and Smith, 2009). In addition, previous mass spectrometry-based studies suggested that hnRNP A1 might contain a PARylation site at K298 and a putative PAR-binding motif (PBM) between the two RRM domains at amino acid (aa) 92–113 (Gagné et al., 2008; Martello et al., 2016). To characterize the PARylation site(s), we generated constructs to express Flag-tag hnRNP A1 of a PARylation site mutant (K298A) or a PBM mutant (R92A-K105/106V, referred to as PBM^mut^ thereafter) in HeLa cells. To examine the significance of PARylation, cells transfected with Flag-tagged wild-type (WT), K298A or PBM^mut^ hnRNP A1 were treated with H_2_O_2_, and the cell lysates were examined by immunoprecipitation (IP) with anti-Flag and Western blotting with anti-PAR. Compared to WT hnRNP A1, PARylation of K298A was dramatically reduced, whereas the association with PAR was not affected in the K298A mutant, evident by the similar levels of co-immunoprecipitation (co-IP) of other PARylated proteins to WT (Fig. 2E). In contrast, PBM^mut^ showed drastically decreased co-immunoprecipitation of other PARylated proteins, whereas its own PARylation was not reduced but unexpectedly increased (Fig. 2E). Of note, hnRNP A1 showed an up-shifting PARylation smear to a similar extent in the *in vitro* PARylation assay (Fig. 2C), indicating that the hnRNP A1 protein was capable of being massively PARylated when induced. Thus, the data suggested that binding to PAR and/or PARylated proteins via the PBM might prevent hyper-PARylation of hnRNP A1 at K298.

### PARylation and PAR-binding are required for hnRNP A1 translocation to SGs

We showed that the cellular PARylation levels affected the recruitment and recovery of hnRNP A1 to and from SGs (Fig. S1 and Fig. 1D-1F). However, it was unclear if PARylation directly regulated the translocation of hnRNP A1 to SGs or it was an indirect effect resulted from altered dynamics of the SGs. To address this question, we expressed the PARylation mutant K298A and PAR-binding deficient PBM^mut^ of hnRNP A1 in HeLa cells and examined their cellular localization before and after stress by immunostaining. In the absence of stress, WT and K298A of hnRNP A1 were predominantly nuclear. PBM^mut^ was mainly localized to the nucleus but also showed cytoplasmic foci that did not co-localize with the SG marker TIAR (Fig. 3A). Of note, the PBM is not located within the RGG or M9 domain, the known nuclear localization sequences of hnRNP A1 (Fig. 2D; Siomi and Dreyfuss, 1995; Nichols et al., 2000). Thus, it is unlikely that the PBM^mut^ cytoplasmic foci are formed due to a defect in nuclear importing of hnRNP A1.

**Figure 3.**
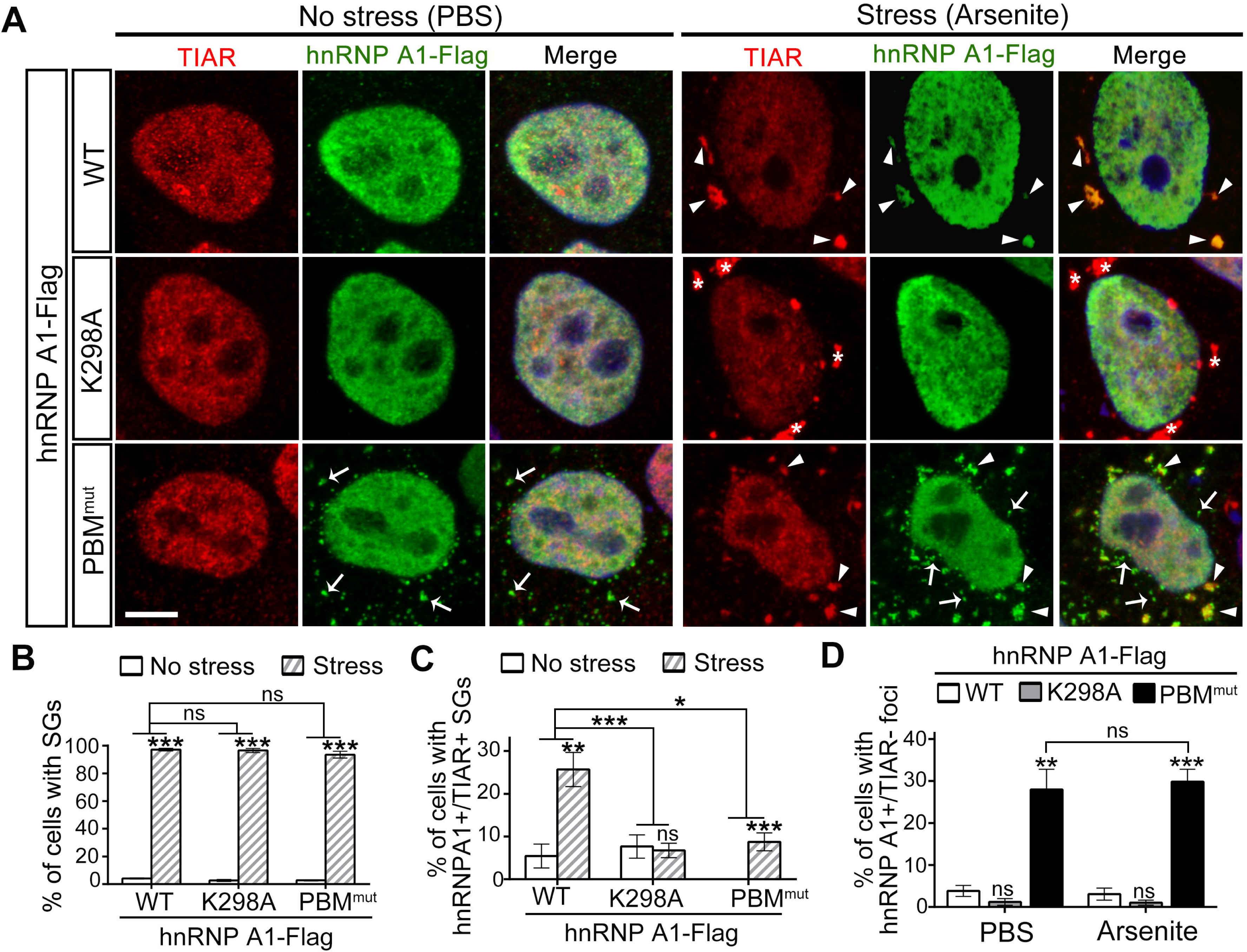
The PARylation and PAR-binding mutants of hnRNP A1 K298A and PBM^mut^ exhibit deficits in translocation and targeting to SGs. (**A**) Representative images of HeLa cells transfected with WT, K298A or PBM^mut^ of Flag-tagged hnRNP A1 and treated with PBS (no stress) or arsenite (stress) for 30 min. Arrows, hnRNP A1+/TIAR-cytoplasmic foci; arrowheads, hnRNP A1+/TIAR+ SGs; asterisks, hnRNP A1-/TIAR+ SGs. (**B-D**) The percentage of cells with SGs (B), hnRNP A1+/TIAR+ SGs (C), or hnRNP A1+/TIAR-cytoplasmic foci (D) is quantified. Mean ± SEM; n = over 300 cells each condition, pooled results of 3 independent repeats; \**p* < 0.05, \*\**p* < 0.01, \*\*\**p* < 0.001; ns, not significant; Student’s *t*-test. Scale bar: 10 μm.

Next, we stressed the cells with arsenite (100 μM, 30 min) which significantly induced the formation of SGs in both WT and the two mutant hnRNP A1 K298A and PBM^mut^ (Fig. 3A). Thus, the overall ability of the cells to form SGs was not affected by expressing K298A or PBM^mut^ hnRNP A1 (Fig. 3B). In response to stress, WT hnRNPA1 showed a significant cytoplasmic translocation and co-localization with TIAR-labeled SGs (Fig. 3A and 3C). Interestingly, K298A mutant was not recruited into SGs and remained predominantly in the nucleus (Fig. 3A and 3C). In response to stress, PBM^mut^ was recruited into TIAR-labeled SGs (Fig. 3A), but the induction was to a less extent than WT hnRNP A1 (Fig. 3C). In addition, the percentage of cells with “PBM^mut^+/TIAR-” cytoplasmic foci showed no difference before and after stress (Fig. 3A and 3D), indicating that stress did not affect the formation or turnover of the PBM^mut^ cytoplasmic foci. Together, these data suggest that PARylation of hnRNP A1 is required for its cytoplasmic translocation, whereas binding to PAR or PARylated proteins regulates the sorting and/or delivery of hnRNP A1 to SGs.

### PAR promotes phase separation of hnRNP A1

RNPs such as hnRNP A1 can phase separate *in vitro*, which may regulate the assembly of RNP granules and the pathological progression to amyloid aggregations *in vivo* (Lin et al., 2015). The findings that hnRNP A1 was not only PARylated but also associated with PARylated proteins *in vitro* and *in vivo* (Fig. 2) prompted us to test whether PAR could directly regulate the LLPS of hnRNP A1. As previously reported (Lin et al., 2015), recombinant hnRNP A1 formed dynamic liquid droplets (LDs) *in vitro*, which increased in size with decreasing concentration of the salt (NaCl, 25–300 mM) and increasing concentration of hnRNP A1 (10–60 μM) (Fig. 4A).

**Figure 4.**
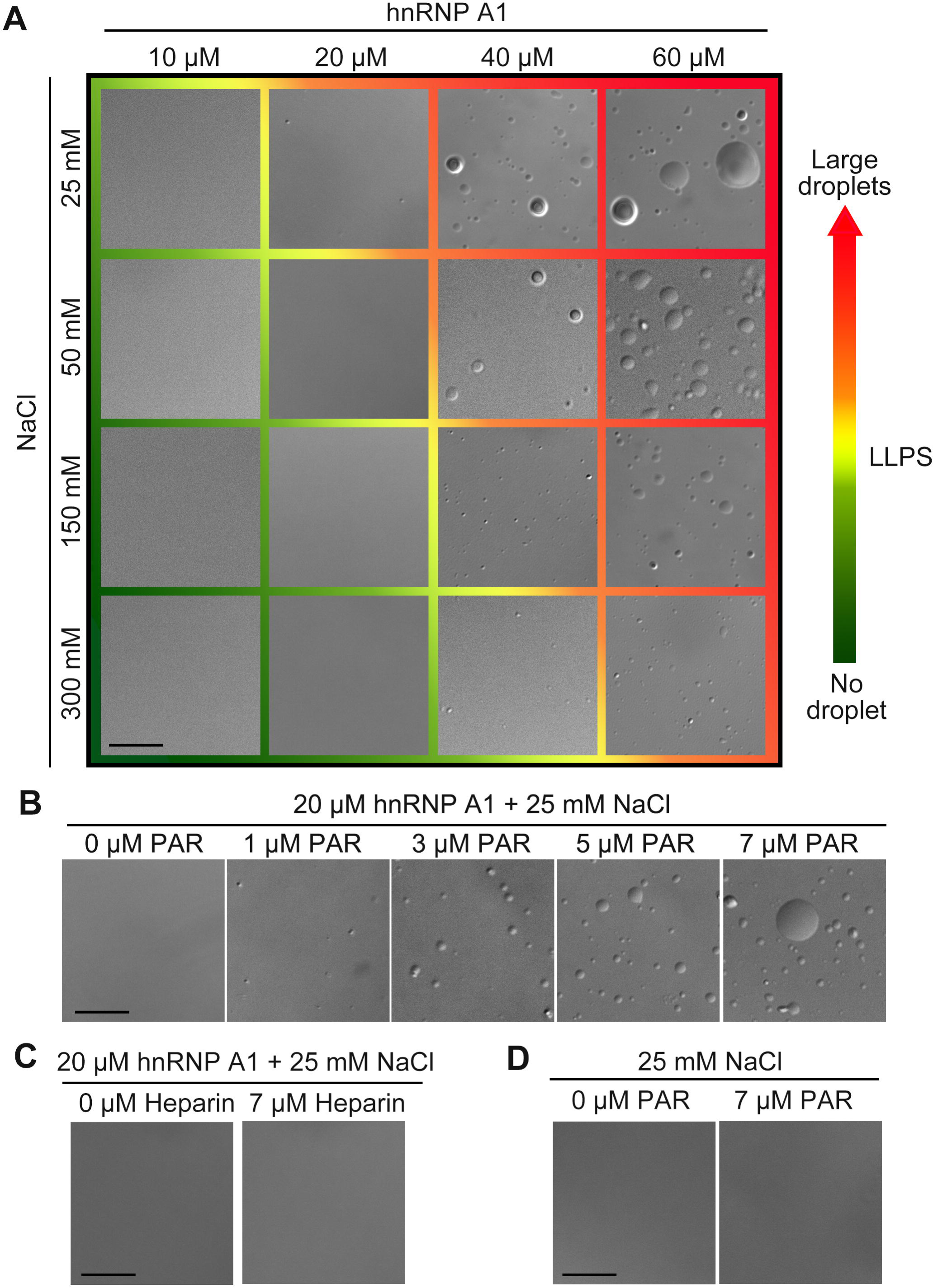
PAR promotes LLPS of hnRNP A1 *in vitro*. (**A**) hnRNP A1 forms dynamic LDs by LLPS *in vitro*. The concentrations of hnRNP A1 and the NaCl are shown. The higher hnRNP A1 and lower NaCl concentrations, the more and larger LDs are formed. (**B**) Addition of PAR promotes LLPS of hnRNP A1 in a dose-dependent manner. (**C**) The same concentration of heparin (7 μM) cannot induce LLPS of hnRNP A1 at the same condition. (**D**) PAR (7 μM) alone does not phase separate *in vitro*. Scale bars: 10μm.

Next, to determine if PAR affected the LLPS of hnRNP A1, we chose a condition at which spontaneous LLPS of hnRNP A1 barely occurred (20 μM hnRNPA1 and 25 mM NaCl, Fig. 4A). We added purified PAR polymers with increasing concentrations (1–7 μM) into the above *in vitro* demixing system, which induced the LLPS of hnRNP A1 in a dose-dependent manner (Fig. 4B). As a control, addition of heparin (7 μM) did not promote the LLPS of hnRNP A1 (Fig. 4C). PAR (7 μM) alone did not alter the phase separate *in vitro* at the same condition (Fig. 4D). Thus, it is a specific effect of PAR to promote LLPS of hnRNP A1.

### hnRNP A1 and TDP-43 co-phase separate *in vitro* and PAR promotes the co-LLPS

A recent study showed that hnRNP A2, another hnRNP family protein, could co-phase separate with TDP-43 and induced co-aggregation (Ryan et al., 2018). Therefore, we examined whether hnRNP A1 and TDP-43 could co-phase separate *in vitro* and whether PAR could influence this process. As shown in Figure 5A, at the condition of 50 μM of hnRNP A1 and 100 mM of NaCl, hnRNP A1 did not phase separate spontaneously. TDP-43^1-274^ (50 μM) but not BSA (50 μM) showed mild LLPS at this condition, as only a few small LDs were spotted (Fig. 5B’-5C’). Mixing hnRNP A1 with TDP-43^1-274^ (50 μM) but not BSA (50 μM) resulted in the formation of massive and much larger LDs (Fig. 5B-5C), suggesting that interaction with each other promoted the co-LLPS of hnRNP A1 and TDP-43^1-274^. However, although TDP-43^274-414^ is more prone to phase separate even at lower concentration (20 μM; Fig. 5D’-5E’), the addition of TDP-43^274-414^ did not promote phase separation of hnRNPA1 (Fig. 5D-5E). To further confirm the increase of LDs when hnRNP A1 and TDP-43^1-274^ were mixed were indeed due to co-LLPS of two proteins rather than a single protein, we prepared fluorophore-labeled hnRNP A1 and TDP-43^1-274^. Similar to the unlabeled proteins, hnRNP A1-Alexa 647 alone did not form any LDs; TDP-43^1-274^-Alexa 555 formed a few small, green only LDs; they together formed large LDs that contained both hnRNP A1 and TDP-43^1-274^ (Fig. 5F). Thus, our results showed that hnRNPA1 co-phase separates with TDP-43^1-274^ *in vitro*.

**Figure 5.**
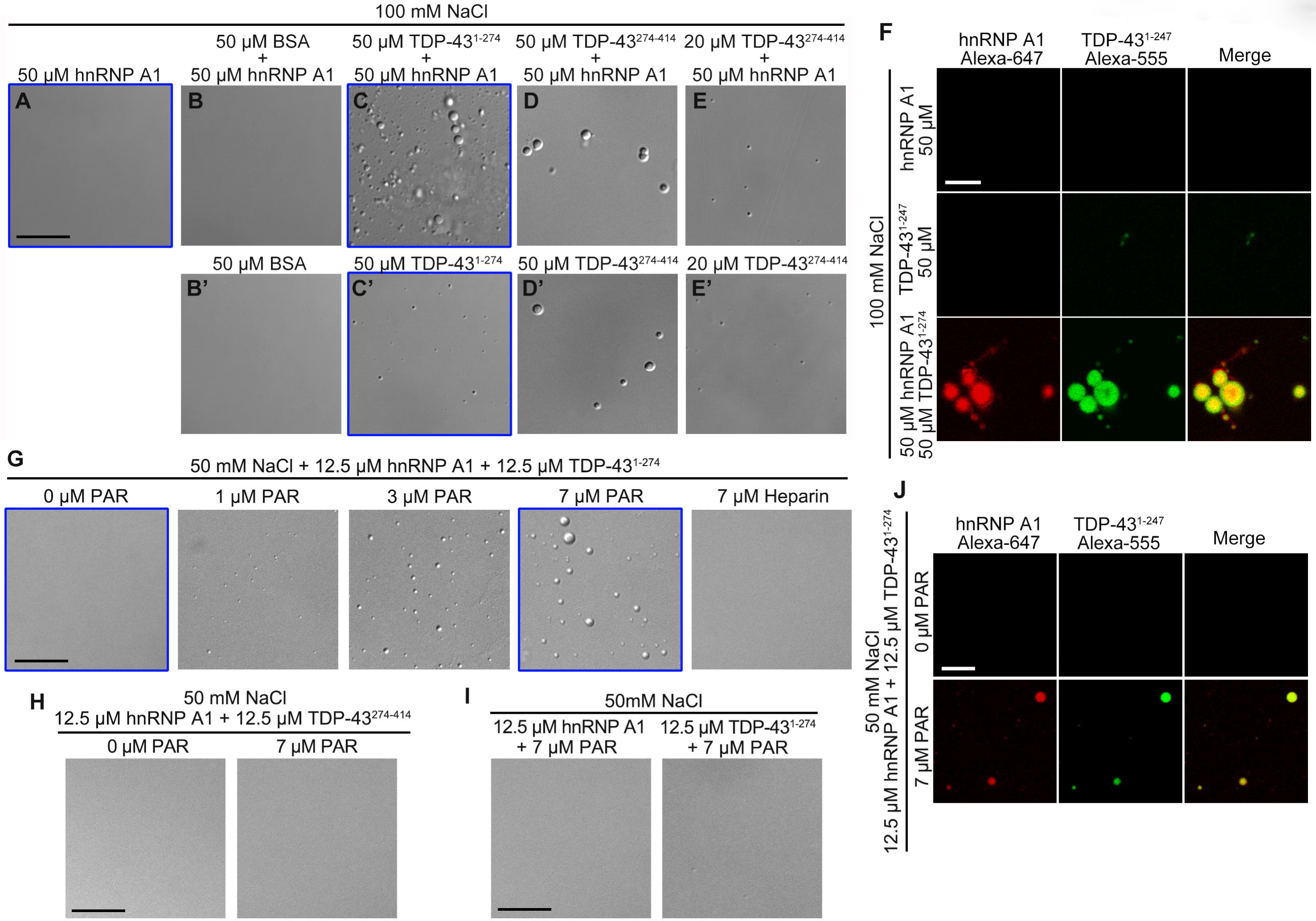
hnRNP A1 and TDP-43 co-phase separate *in vitro* and PAR affects this process. (**A)** hnRNP A1 alone does not form spontaneous LDs by LLPS at the indicated condition *in vitro*. (**B**-**E**) co-LLPS experiments of hnRNP A1 with BSA (B), TDP-43^1-274^ (C), or TDP-43^274-414^ (D-E) with the indicated concentrations of each protein. (**B’**-**E’**) LLPS experiments of BSA (B’), TDP-43^1-274^ (C’), or TDP-43^274-414^ (D’-E’) alone at the same condition as in (B-E). (**F**) Fluorphore-conjunct hnRNP A1 (AlexaFluor-647, red) and TDP-43^1-247^ (AlexaFluor-555, green) confirms that the LDs formed by co-LLPS contain both of the two proteins. All tests in A-F are carried out with 100 mM of NaCl. (**G**) PAR but not heparin promotes the co-LLPS of hnRNP A1 and TDP-43^1-274^ *in vitro*. (**H**) PAR cannot induce the co-LLPS of hnRNP A1 and TDP-43^274-414^. (**I**) At the same condition of (G), PAR does not trigger the LLPS of only hnRNP A1 or only TDP-43^1-274^. (**J**) In the presence of PAR (7 μM), co-LLPS of labeled hnRNP A1 (AlexaFluor-647, red) and TDP-43^1-247^ (AlexaFluor-555, green) occurs so that the LDs contain both red and green fluorescence. The conditions highlighted by a blue box in A-E’ and G are examined in F and J, respectively. The concentration of each component in the demixing system is indicated. Fluorescent images are shown in pseudo colors. Scale bars: 5 μm in F and J, all others 10 μm,

We then examined whether PAR could affect the co-LLPS of hnRNP A1 and TDP-43. To test this, we lowered the concentration of hnRNP A1 and TDP-43^1-274^ to 12.5 μM each and NaCl to 50 mM, at which spontaneous co-LLPS did not occur (Fig. 5G). Addition of PAR (1–7 μM) but not heparin (7 μM) promoted co-LLPS of hnRNP A1 and TDP-43^1-274^ in a dose-dependent manner (Fig. 5G). Again, we did not observe co-LLPS of hnRNP A1 with TDP-43^274-414^ or any effect of PAR (7 μM) on their co-LLPS (Fig. 5H). Of note, no LLPS was triggered when PAR (7 μM) was added to hnRNP A1 (12.5 μM) or TDP-43^1-274^ (12.5 μM) alone (Fig. 5I). Also, the fluorophore-labelel hnRNP A1 and TDP-43^1-274^ confirmed that the LDs triggered by addition of PAR in Figure 5G contained both of the two proteins that co-phase separated *in vitro*. (Fig. 5J).

### PARylation modulates the interaction between hnRNP A1 and TDP-43

Next, to investigate whether hnRNP A1 interacted with TDP-43 *in vivo* and how PARylation regulated this process, we conducted the co-IP experiments. We found that the endogenous TDP-43 protein could be co-immunoprecipitated with transiently expressed hnRNP A1-Flag, which was consistent with the observations that TDP-43 bound to a few hnRNP family proteins (Buratti et al., 2005; D’Ambrogio et al., 2009). Further, we showed that the interaction between hnRNP A1 and TDP-43 depended on the presence of RNA (Fig. S4). Next, we tested the effect of the PARP inhibitor Olaparib, which markedly reduced the amount of endogenous TDP-43 that were co-immunoprecipitated with hnRNP A1; on the other hand, activation of PARP1 by H_2_O_2_ moderately increased the co-IP of TDP-43 with hnRNP A1 (Fig. 6A-6B).

**Figure 6.**
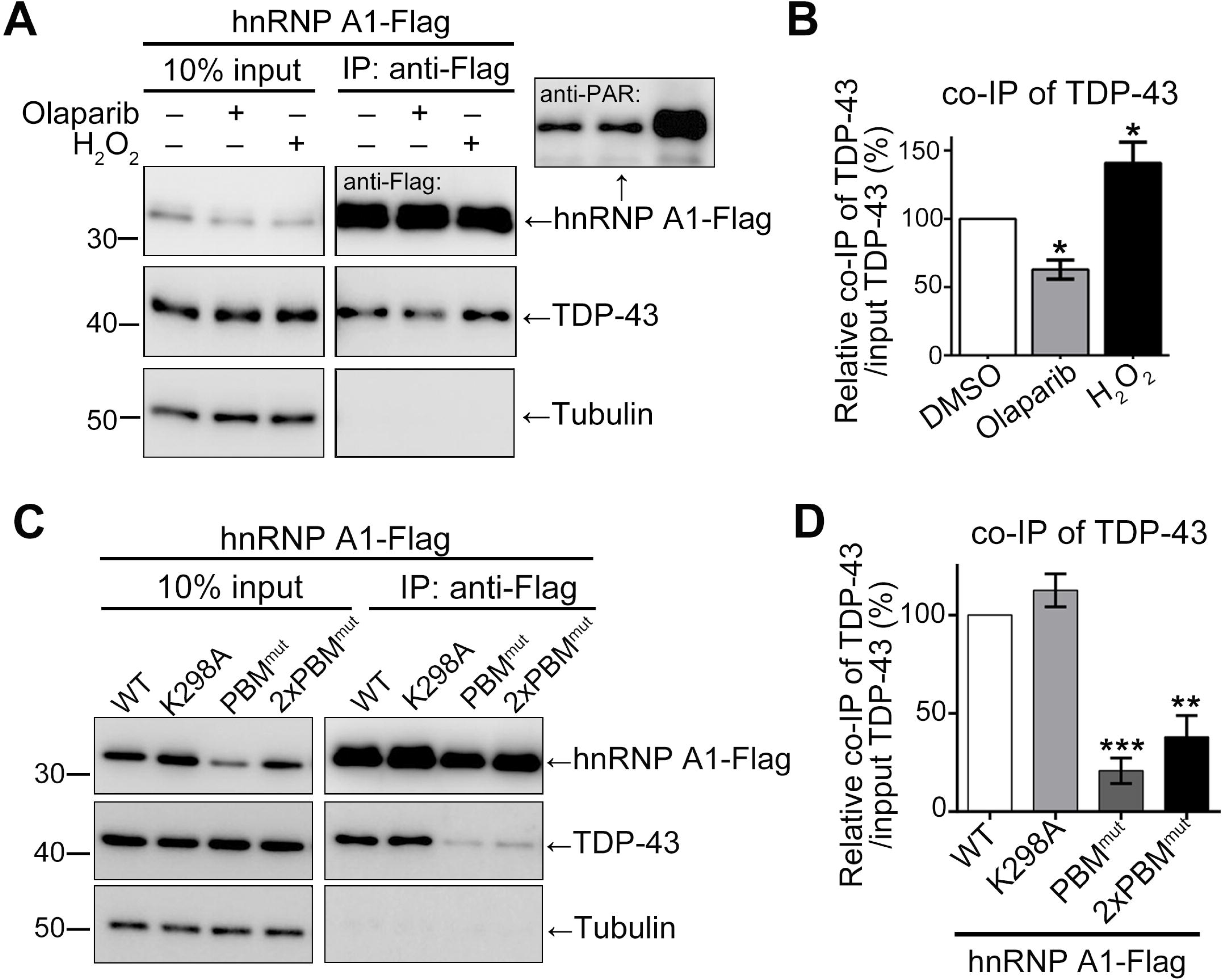
PARrlation modulates the interaction between hnRNP A1 and TDP-43. (**A**) Flag-tagged hnRNP A1 is expressed in HeLa cells treated with vehicle control (DMSO), Olaparib (20 μM) or H_2_O_2_ (500 μM), and immunoprecipitated with anti-Flag. The immunoprecipitates are then separated by SDS-PAGE and examined by Western blotting. (**B**) The level of co-IP of TDP-43 with hnRNP A1 is quantified as ratio of co-IP of TDP-43/input TDP-43 and shown as percentage to the DMSO control group. (**C**) Western blot analysis of cells transfected with WT, K298A or PBM^mut^ of Flag-tagged hnRNP A1, immunoprecipitated with anti-Flag, and blotted with indicated antibodies. 2xPBM^mut^, cells transfected with 2 times of the PBM^mut^ expression plasmids. (**D**) The relative level of TDP-43 co-immunoprecipitated with WT or mutant hnRNP A1 is normalized to TDP-43 level in the input and shown as percentage to the WT hnRNP A1 group. Means ± SEM, n = 5 (B) or 3 (D); *p < 0.05, \*\**p* < 0.01, \*\*\**p* < 0.001; One-way ANOVA.

To understand how PARylation and PAR-binding regulated the interaction of hnRNP A1 with TDP-43, we examined the binding affinity of WT, K298A and PBM^mut^ with TDP-43 by co-IP. Compared to WT hnRNP A1, the PARylation site mutant K298A showed a similar capability of pull-down of TDP-43, however, the association of TDP-43 with PBM^mut^ was dramatically reduced (Fig. 6C-6D). Of note, the protein level of PBM^mut^ in the input was lower due to reduced solubility (Fig. S5). To be able to compare the co-IP efficiency with a similar input level, cells transfected with 2 times of the PBM^mut^ expression plasmids (2xPBM^mut^) were also examined. The input protein levels of PBM^mut^ were significantly improved, however, co-IP of TDP-43 was still barely detected (Fig. 6C-6D). Together, these data indicate that the association of PAR or PARylated proteins via the PBM of hnRNP A1 is required for the interaction between hnRNP A1 and TDP-43.

### Inhibition of PARylation reduces the cytotoxicity of hnRNP A1 and TDP-43 in motor neuron-like NSC-34 cells

We extended the study of the impact of PARylation on biochemical properties to functional readouts such as the cytotoxicity of these disease-associated RNPs. We used the mouse motor neuron (MN)-like hybrid cell line, NSC-34 cells, to overexpress hnRNP A1 or TDP-43 by lentiviral infection. The NSC-34 cells overexpressing hnRNP A1 or TDP-43 OE cells exhibited remarkable morphological changes and significant reduction of cell viability (Fig. 7A-7B and Fig. S6), which confirmed the cytotoxicity of overexpression (OE) of hnRNP A1 or TDP-43 in the MN-like cell model.

**Figure 7.**
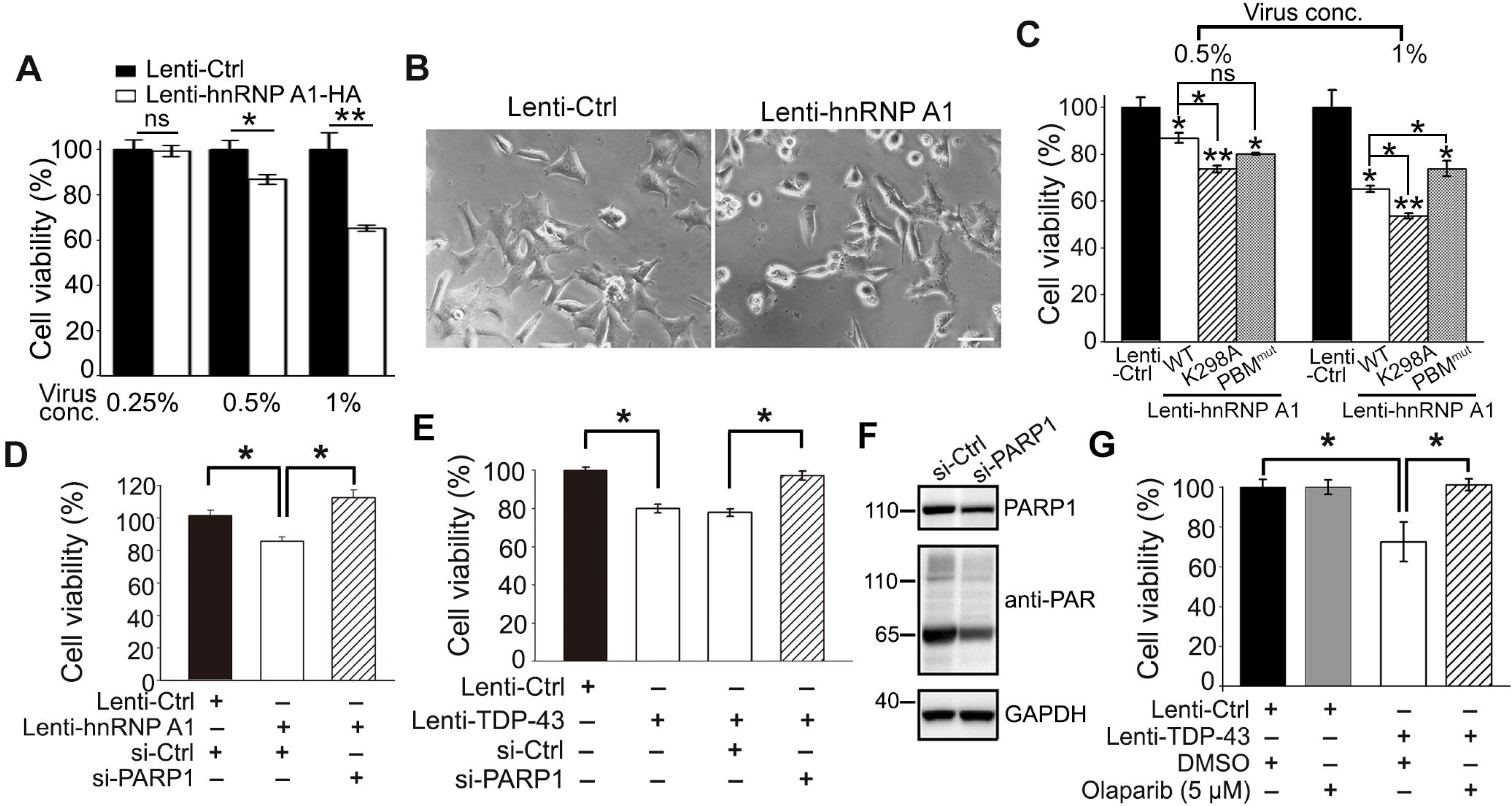
PARP inhibition mitigates the cytotoxicity of hnRNP A1 and TDP-43 in MN-like NSC-34 cells. (**A**) NSC-34 cells infected with increased concentrations of lenti-hnRNP A1-HA show decreased cell viability in the Cell Counting Kit-8 (CCK-8) assay, indicating that hnRNP A1-induced cytotoxicity is dose-dependent. (**B**) Representative bright field images of NSC-34 cells infected with Lenti-Ctrl or Lenti-hnRNP A1-HA (1% virus concentration) are shown. Scale bar: 50 μm. (**C**) WT, K298A and PBM^mut^ of hnRNP A1 exhibit different levels of cytotoxicity in NSC-34 cells. (**D-E**) NSC-34 cells overexpressing hnRNP A1 (C) or TDP-43 (D) by lentivirus infection show decreased cell viability in the CCK-8 assay, which is significantly suppressed by KD of PARP1 by (si-PARP1) compared to the scrambled siRNA control (si-Ctrl). (**F**) Western blot analysis confirms PARP1 KD and decrease of overall PARylation levels by siRNA of PARP1 (si-PARP1). (**G**) PARP inhibitor Olaparib rescues TDP-43 OE-induced reduction of cell viability in the NSC-34 model. Mean ± SEM; n = 3; \**p* < 0.05, \*\**p* < 0.01; ns, not significant; Student’s *t*-test.

The K298A and PBM^mut^ mutants exhibited different levels of cytotoxicity compared to WT hnRNP A1 (Fig. 7C). The K298A mutant was more toxic than WT hnRNP A1 at both low and high concentrations of lentiviral infection. Considering that PARylation at K298A was required for cytoplasmic translocation of hnRNP A1 to SGs (Fig. 3), this result suggests that the inability of promptly responding to cellular stress enhances the cytotoxicity of disease-associated RNPs. PBM^mut^ showed a similar level of cytotoxicity to WT at a low concentration of lentiviral infection. At a higher infection concentration, the cytotoxicity of WT and K298A increased drastically, but that of PBM^mut^ did not change much and hence became less toxic than WT and K298A at this condition (Fig. 7C). It was possible that the decreased binding to PARylated proteins made PBM^mut^ and the SGs it was associated with more dynamic and less solid than those of WT hnRNP A1. This would be especially important when the proteins were expressed at high levels, as it may reduce the chance of the SGs to develop into gel-like or solid protein aggregations that resemble the disease pathology.

As K298A and PBM^mut^ affected the cytotoxicity of hnRNP A1 differently, we were keen to know the overall consequence of reducing PARylation levels in cells and whether it could mitigate the cytotoxicity of hnRNP A1 or TDP-43. Thus, we treated the NSC-34 cells with siRNA of PARP1 (si-PARP1). We found that downregulation of PARP1 significantly suppressed the decrease of cell viability induced by hnRNP A1 or TDP-43 OE (Fig. 7D-7F). Furthermore, as an attempt to testing small-molecule for treating ALS, we examined the PARP inhibitor Olaparib. We tested different doses and found that 5 μM of Olaparib did not reduce the viability of NSC-34 cells but showed a remarkable suppression of TDP-43 OE-mediated cytotoxicity (Fig. 7G). Thus, both genetic and pharmacological inhibition of PARP significantly suppressed the cytotoxicity of ALS-associated RNPs in MN-like NSC-34 cells.

### Downregulation of *Parp* suppresses TDP-43-mediated neurodegeneration in a *Drosophila* model of ALS

Finally, we validated these findings in an *in vivo* model of ALS using transgenic flies expressing human TDP-43 (hTDP-43). hTDP-43 OE in the fly photoreceptor cells (GMR driver) caused age-dependent eye degeneration, which was drastically suppressed by transgenic downregulation of fly *Parp* (RNAi-*Parp*) compared to RNAi-Ctrl (Fig. 8A-8B). We examined the KD efficiency of RNAi-*Parp* by quantitative real-time PCR (qPCR) assay and the mRNA level of *Parp* was reduced to below 40% of the RNAi-ctrl group (Fig. S7A). Also, we showed that the reduction of *Parp* did not affect the protein levels or solubility of hTDP-43 (Fig. S7B-S7C), confirming that the suppression by RNAi-*Parp* was not due to a reduction of transgenically expressed hTDP-43 protein in the system. Furthermore, we performed the climbing and lifespan assays to evaluate the behavioral consequences, which might represent more closely to the disease-relevant symptoms in ALS. We induced hTDP-43 OE in adult fly neurons using an *elav*GS driver and added RU486 to the fly food (80 µg/ml) starting from day one of the adulthood. It caused an age-dependent decline of the climbing capability and a significant shortening of the lifespan, both of which could be suppressed by RNAi-*Parp* in the fly neurons (Fig. 8C-8D). Together, these data indicate that PARP can modify TDP-43-mediated neurodegeneration *in vivo*.

**Figure 8.**
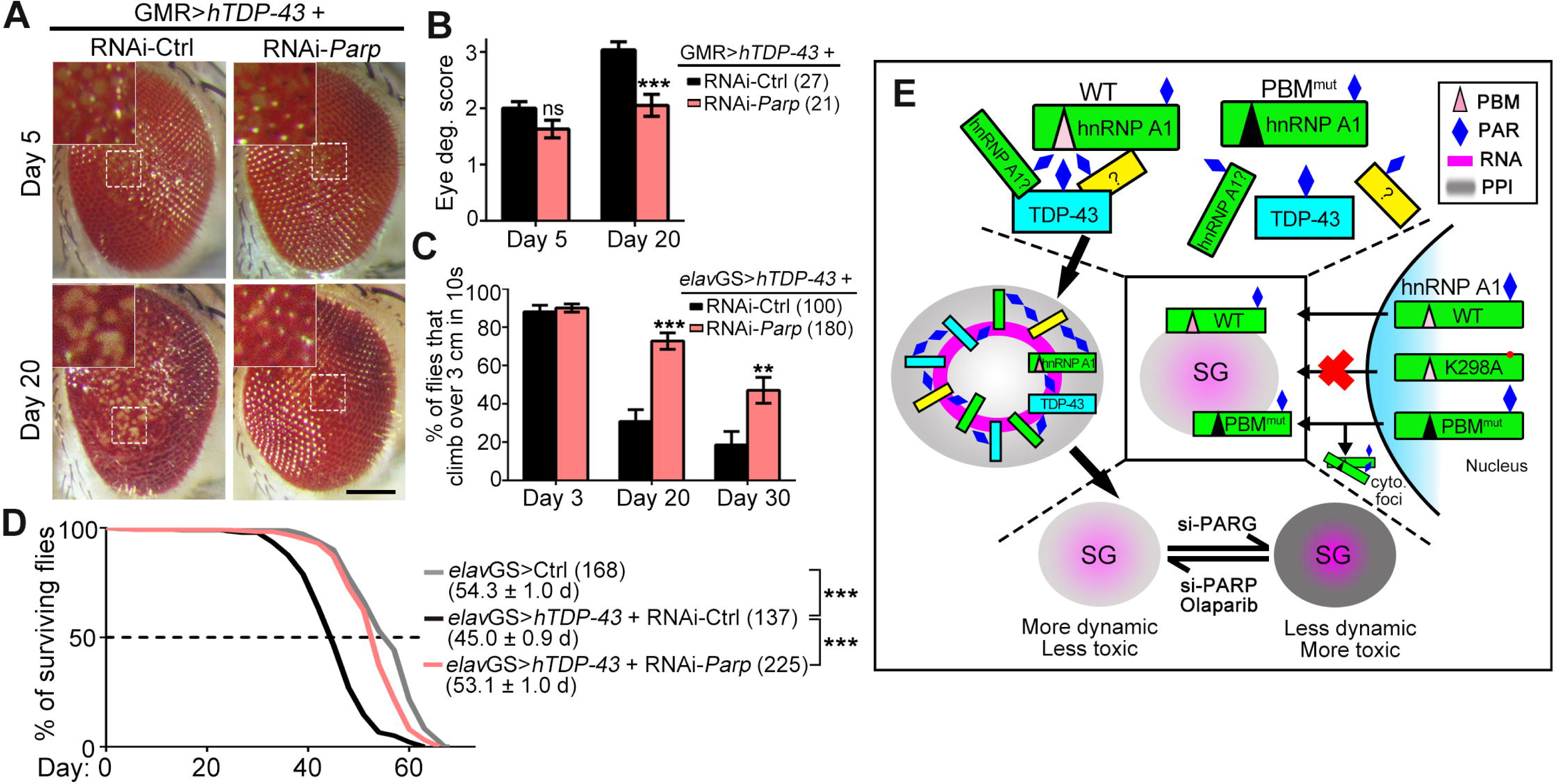
Downregulation of *Parp* alleviates neurodegeneration in a *Drosophila* model of ALS. (**A-B**) Representative images (A) and quantification of the degeneration scores (B) of the fly eyes (GMR-Gal4) expressing hTDP-43 with RNAi-*Parp* or RNAi-Ctrl (RNAi-*mCherry*). (**C-D)** RNAi KD of *Parp* in adult fly neurons using an *elav*GS driver (induced with RU486, 80 µg/ml, starting from day one of the adulthood) alleviates TDP-43-mediated climbing decline (C) and shortened lifespan (D). The climbing capability is evaluated as the average percentage of flies that climb over 3 cm within 10 seconds. The median lifespan and the number of flies tested for each genotype are indicated. Ctrl, UAS-*Luciferase*; RNAi-Ctrl, UAS-RNAi-*mCherry*. Mean ± SEM; \*\**p* < 0.01, \*\*\**p* < 0.001; ns, not significant; Student’s *t*-test. Scale bar, 100 μm. (**E**) A schematic model of the role of PARylation in regulating SGs and disease-related RNPs. hnRNP A1 is PARylated at K298 and binds to PAR and/or PARylated proteins via the PBM. PARylation and PAR-binding regulate the recruitment of hnRNP A1 to SGs: K298A mutant does not respond to stress and remains predominantly nuclear; PBM^mut^ can be recruited to SGs, but a portion of it forms mis-localized cytoplasmic foci that do not co-localize with SGs. The mis-localization of PBM^mut^ is likely due to reduced interaction with other RNPs such as TDP-43. The association and assembly of RNP granules require RNAs, while PARylation modulates the protein-protein interaction between RNPs. Thus, PARylation may act as a molecular glue to promote LLPS and enhance the assembly of SGs by cross-linking different RNPs. Pharmacological or genetic inhibition of PARP improves the dynamics of RNP granules and mitigates the neurotoxicity of hnRNP A1 and TDP-43. PAR, poly(ADP-ribose); PBM, PAR-binding motif; PPI, protein-protein interaction; WT, wild-type hnRNP A1; K298A, PARylation site mutant of hnRNP A1; PBM^mut^, PAR-binding motif mutant of hnRNP A1; SG, stress granule; cyto. foci, cytoplasmic PBM^mut^ foci.

## DISCUSSION

### PARylation regulates SG dynamics and phase separation of RNPs

In this study, we reveal that decrease of the cellular PARylation levels suppresses the formation of SGs and the recruitment of ALS-related RNPs such as TDP-43 and hnRNPA1 to SGs, while increase of PARylation levels delays the disassembly of SGs and the recovery of the RNPs. Interestingly, the function of PARylation appears to be specifically enriched in regulation of RNA/DNA-binding proteins and the associated complexes. For example, PARylation is involved in the regulation of the mitotic spindle (Chang et al., 2004), Cajal bodies (Kotova et al., 2009), DNA damage repair (Ahel et al., 2009), and microRNA-mediated translational repression (Leung et al., 2011). Because of their essential role in RNA processing and homeostasis (Dreyfuss et al., 1993; Lee et al., 2012), dysregulation of RNP granules contributes to the pathogenesis of neurodegenerative diseases (Anderson and Kedersha, 2009; Buchan and Parker, 2009; Zhang et al., 2018). This is at least in part because many of the RNPs contain intrinsically disordered LC domains that can undergo spontaneous self-assembly via LLPS to generate higher-order structures such as solidified LDs, and irreversible amyloid fibrils. We find that adding PAR to the demixing system dramatically promotes the LLPS of hnRNP A1 as well as the co-LLPS of hnRNP A1 with TDP-43. Consistently, during manuscript preparation of this study, an independent study recently published online reporting that the liquid demixing of TDP-43 could be affected by PAR (McGurk et al., 2018). Thus, the reversible modification of PARylation may serve as an important regulator of the dynamics of RNP granules, which in case of lasting stimuli or excessive PARylation, pathological irreversible RNP aggregates may form (Fig. 8E).

### The dual regulations of hnRNP A1 by PARylation and PAR-binding

hnRNP A1 can form protein complexes with TDP-43 and other RNPs to regulate RNA processing (Buratti et al., 2005; Mohagheghi et al., 2016). Indeed, we find that hnRNP A1 and TDP-43 can co-phase separate *in vitro* and PARylation strongly regulates their interaction in *vitro* and in cells. The hnRNP family proteins constitute an important component of the SGs and are highly regulated at molecular and cellular levels. For example, pathogenic mutations in the GRD of hnRNP A1 accelerate its recruitment to SGs and the fibrillization (Kim et al., 2013), while methylation of hnRNP A2 in the RGG domain suppresses the phase separation of hnRNP A2 (Ryan et al., 2018).

In this study, we show that PARylation regulates the LLPS of hnRNP A1 *in vitro* and the translocation of hnRNP A1 to SGs *in vivo*. We further reveal that hnRNP A1 contains a PARylation site at K298 that is in the GRD as well as a PBM that resides in between the two RRMs. K298 localizes at the C-terminus of the M9 domain, a non-classical nuclear localization sequence of hnRNP A1. Of note, K298 is not within the M9 core sequence (SNFGPMKGGNFGGRSSGPY) that is crucial for the nuclear import and binding to transportin (Iijima et al., 2006). Indeed, the PARylation deficient K298A mutant shows no defect in nuclear importing. Instead, it does not translocate to the cytoplasm in response to stress and exhibits the strongest toxicity among the three hnRNP A1 species tested in this study (see Fig. 7). These data implicate the PARylation at K298 as an important mechanism for hnRNP A1 to sense stress and/or serve as a nuclear exporting signal.

The PAR-binding deficient PBM^mut^ of hnRNP A1 can respond to stress and translocate to SGs, likely because the signaling mechanims by PARylatino at K298 is intact. However, the SG recruitment of PBM^mut^ is less effective than WT and a portion of PBM^mut^ forms mis-localized cytoplasmic foci that do not co-localize to SGs. Of note, PBM^mut^ shows hyper-PARylation in the *in vitro* assay (see Fig. 2C), possibly at K298 of hnRNP A1, which may result in abnormal activation of the nuclear exporting of PBM^mut^. In the meanwhile, since PBM^mut^ shows a greatly reduced capacity to pull down PARylated proteins, we speculate that the association with PAR or PARylated proteins ensures the proper transport and anchoring of hnRNP A1 with SGs. Despite of forming cytoplasmic foci, PBM^mut^ is not more toxic than WT hnRNP A1. Instead, PBM^mut^ is less toxic especially when expressed at high levels (see Fig. 7). It is possibly because the reduced association of PBM^mut^ with SGs makes the granules less prone to develop into pathogenic aggregates. And this effect may be particular important when the protein is at high abundance as LLPS of hnRNP A1 occurs with increasing concentrations and PAR promotes this process in a dose-dependent manner. It is worth noting that this is unlike the PBM mutant of TDP-43, which is thought to be more toxic because it could not be recruited to SGs and formed hyper-phosphorylated cytoplasmic foci (McGurk et al., 2018). Nevertheless, downregulation of PARylation levels reduces the cytotoxicity of hnRNP A1 and TDP-43 in both cases (see Fig. 7), which suggests that, under disease conditions, reduction of PAR-mediated association of RNPs and restore of the dynamics of RNP granules may be beneficial.

### Potential therapeutic values of PARP inhibitors for neurodegenerative diseases

Small-molecule inhibitors of PARP enzymes have received increasing interests since the initial discovery of them in killing *BRCA1/2*-mutant cancer cells. While the involvement of PARP in DNA damage repair has heralded the development of PARP inhibitors in cancer biology and therapy (Davar et al., 2012), the functions of PARP are beyond DNA repair and oncology (Bai, 2015). For example, the neuroprotective effects of PARP inhibitors have been reported in Hungtinton’s disease, cerebral ischemia and axonal regeneration (Cardinale et al., 2015; Egi et al., 2011; Teng et al., 2016; Brochier et al., 2015). In this study, we show that Olaparib, the PARP inhibitor approved by FDA for treating ovarian cancer and breast cancer, can significantly reduce the neurotoxicity of hnRNP A1 and TDP-43 in a motor neuron-like cell line, likely due to the pivotal role of PARP in regulating the protein-protein interaction and the dynamics of the disease-associated, aggregation-prone RNP granules. Therefore, we propose Olaparib as a candidate for developing ALS drugs, and further animal and clinical studies are needed to evaluate the potential therapeutic values of Olaparib and other PARP inhibitors to treat ALS and other RNP-related diseases.

## MATERIALS AND METHODS

### Plasmids and constructs

To generate pCAG-TDP-43-HA, pCMV-TDP-43-3xFlag and pBID-UASC-TDP-43 plasmids, human TDP-43 DNA was amplified from a TDP-43-Myc plasmid (Jiang et al., 2013) by PCR using the primers specified below. The desired PCR products were subcloned into a pCAG vector (Chen et al., 2014) or a pCMV-3Tag-3B vector (Agilent Technologies) using the ClonExpress™ II One Step Cloning Kit (Vazyme). To generate pCAG-hnRNP A1-Flag plasmid, human hnRNP A1 coding sequence was amplified from the cDNA of HeLa cells using by PCR and sub-cloned into a pCAG vector by homologous recombination using the ClonExpress™ II One Step Cloning Kit (Vazyme). The pCAG-K298A-Flag and pCAG-PBM^mut^-Flag of hnRNP A1 were generated by PCR using the pCAG-hnRNP A1-Flag as a template and hnRNP A1 by site-directed mutagenesis using the Fast Mutagenesis Kit II (Vazyme).

For *Escherichia coli* (*E. coli*) expression, the pET-28a-TDP-43^1-274^-6xHis and pET-28a-TDP-43^274-414^-6xHis constructs were generated by PCR amplification of the truncated TDP-43 fragments from the above full-length TDP-43 plasmid and inserted between the BamHI and Xho1 sites in pET-28a-6*his vector. The pET9d-hnRNP A1 plasmid was obtained from Addgene (Plasmid #23026). To generate lentiviral packaging constructs to express Flag, TDP-43-Flag or hnRNP A1-HA in NSC-34 cells, the coding sequence was amplified by PCR using the above expression plasmids as the template and each of the PCR products was sub-cloned into the pCDH-CMV-MCS-EF1-Puro vector using the restriction enzyme sites including EcoRI, Xbal, Xhol, BstBI and BamHI.

Primers used for PCR to generate the expression plasmids are summarized below. All constructs were verified by sequencing to ensure the integrity of the cloned open reading frames.

**Table 1.**
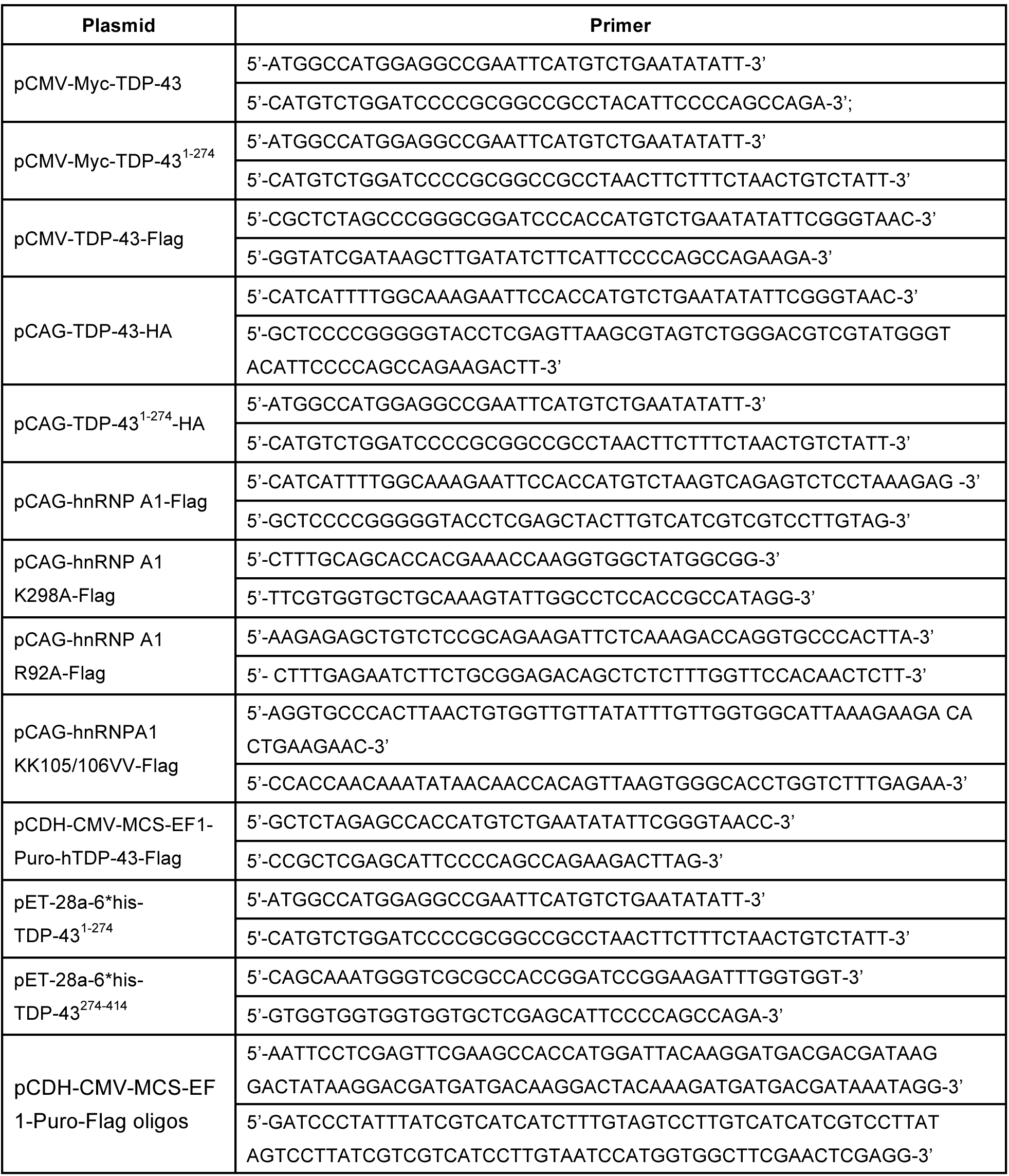
Primers and oligos used to generate expression constructs in this study.

### Cell culture and transfection

Both HeLa and HEK-293T cells were cultured in Dulbecco’s modified Eagle’s medium (sigma, D0819) supplemented with 10% fetal bovine serum (FBS, BioWest). NSC-34 cells were cultured in RPMI 1640 medium (Gibco, 11875-093) containing 10% (v/v) FBS. All cells were incubated at 37°C in a humidified atmosphere of 95% air and 5% CO_2_. PolyJet™ reagent (SignaGen, SL100688) was used for transfection of plasmids into HeLa and HEK-293 T cells. Cells were transfected for 24–72 h before proceeding with subsequent experiments. For knockdown experiments, siRNA (Genepharma, Shanghai) were transfected into cells using Lipofectamine™ RNAiMAX Transfection Reagent (Invitrogen, 13778150) according to the manufacture’s instruction. siRNA was incubated for 48-72 h before cells were harvest. The siRNA oligos used in this study are listed below:

si-Ctrl: 5’-GCGGUGAAGUUAGAUUACATT-3’

si-hPARG: 5’-GCGGUGAAGUUAGAUUACATT-3’

si-mPARP1: 5’-CGACGCUUAUUACUGUACUTT-3’

### Stress granule induction

HeLa cells grown on coverslips in a 12-well plate were treated with 100 µM of NaAsO_2_ or PBS for 30 min before they were fixed with 4% paraformaldehyde. For recovery, the medium containing NaAsO2 was removed and the cells were incubated in fresh medium for indicated time before fixation. Formation of stress granules was evaluated by subsequent immunocytochemistry assays.

### Immunocytochemistry and confocal imaging

After transfection and drug treatments, cells grown on the cover slips in 24-well plate were fixed with 4% paraformaldehyde in PBS for 15 min at room temperature, permeabilized in 0.5% Triton X-100 in PBS for 10 min, and blocked with 3% goat serum in PBST (PBS + 0.1% Triton X-100) for 30 min at room temperature. The primary and secondary antibodies in the blocking buffer were then incubated at 4°C overnight or at room temperature for 1 h. After 3 washes with PBST, cells were mounted on glass slides using Vectashield Antifade Mounting Medium with DAPI (Vector Laboratories). Fluorescent images were taken with Leica TCS SP8 confocal microscopy system using a 100X oil objective (NA=1.4). Images were processed and assembled into figures using LAS X (Leica) and Adobe Photoshop CS6.

### Protein extraction and Western blotting

Total protein was extracted from cells in 2% SDS extraction buffer (50 mM Tris pH 6.8, 2% SDS, 1% mercaptoethanol, 12.5% glycerol and 0.04% bromophenol blue) containing the protease inhibitor cocktail (Roche, 04693132001), 20 µM Olaparib (Selleck, S1060) and 8 µM ADP-HPD (Millopore, 118415). To separation of soluble and insoluble proteins, cells or fly heads were lysed on ice in RIPA buffer (50 mM Tris pH 8.0, 150 mM NaCl, 1% NP-40, 5 mM EDTA, 0.5% sodium deoxycholate, 0.1% SDS) supplemented with protease and phosphatase inhibitors. After sonication, the homogenates were centrifuged at 13,000 x g for 10-20 min at 4°C. The supernatant was used as the soluble fraction and the pellets containing insoluble fractions were dissolved in a 9 M urea buffer (9 M urea, 50 mM Tris buffer, pH 8.0) after wash.

Proteins were separated by 10% Bis-Tris SDS-PAGE (Invitrogen), immunoblotted with the primary and secondary antibodies. Detection was performed using the High-sig ECL Western Blotting Substrate (Tanon). Images were captured using an Amersham Imager 600 (GE Healthcare) and densitometry was measured using ImageQuant TL Software (GE Healthcare). The contrast and brightness were optimized equally using Adobe Photoshop CS6 (Adobe Systems Inc.). All experiments were normalized to GAPDH or tubulin as indicated in the figures.

### Co-immunoprecipitation

HeLa cells were lysed in IP buffer (25 mM Tris•HCl pH 7.4, 150 mM NaCl, 1% NP-40, 1 mM EDTA, 5% glycerol) containing protease inhibitor cocktails, 20 µM Olaparib and 8 µM ADP-HPD. To test if the interaction between hnRNP A1 and TDP-43 was RNA-dependent, cell lysates were treated with 100 mg/ml of RNase A for 30 min (Qiagen), and then incubated with mouse anti-Flag or rabbit anti-HA antibody on a rotary shaker at 4°C overnight. Mouse or rabbit IgG (Santa Cruz, sc-2025 or sc2027) was used as a control for the pull-down specificity. Anti-FLAG^®^ M2 Affinity Gel (Sigma, A2220) or Dynabeads^®^ Protein G beads (Novex) were then added and incubated at room temperature for 2 h. The gel or beads were then collected according to the manufacture’s instruction and eluted in the 2xLDS sample buffer (Invitrogen) before the subsequent Western blotting assay.

### Antibodies

The following antibodies were used for Western blotting, immunoprecipitation and immunocytochemistry assays: mouse anti-FLAG (Sigma, F3165), mouse anti-HA (Proteintech, 66006-1), anti-pan-ADP-ribose binding reagent (Millipore, MABE1016), rabbit anti-HA (CST, 3724T), rabbit anti-TDP-43 (Proteintech, 10782-2-AP), rabbit anti-TIAR (CST, 8509S), rabbit anti-hnRNPA1 (CST, 8443S), rabbit anti-PAR (ENZO, ALX-210-890A-0100), mouse anti-GAPDH (Proteintech, 60004-1) and mouse anti-Tubulin (MBL, PM054). HRP conjugated secondary antibodies, goat anti-mouse (Sigma, A4416) and goat anti-rabbit (Sigma, A9169). Fluorescent secondary antibodies: goat anti-mouse-Alexa Fluor 488 (Life Technologies, A11012); goat anti-rabbit-Alexa Fluor 568 (Life Technologies, A11012).

### RNA extraction and real-time quantitative PCR (qPCR)

Total RNAs were extracted from cells or fly heads (40 heads per group) using TRIzol (Invitrogen) according to the manufacturer’s instructions. After DNase (Promega) treatment to remove genomic DNA, the reverse transcription was performed using the High-Capacity cDNA Reverse Transcription Kit (Applied biosystems). The cDNA was then used in real-time PCR experiment using the SYBR Green qPCR Master Mix (Bimake) with the QuantStudio™ 6 Flex Real-Time PCR system (Life Technologies). The mRNA levels of *actin* were used as an internal control to normalize the mRNA levels of genes of interest. The primers for qPCR are listed below:

*dParp*:

5’-ATGAAGTACGGAGGCCAACC-3’; 5’-TCTTCACCTGACGCAAACCA-3’

*dActin*:

5’-GAGCGCGGTTACTCTTTCAC-3’; 5’-GCCATCTCCTGCTCAAAGTC-3’

### Purification of TDP-43 and hnRNP A1

TDP-43^1-274^: TDP-43^1-274^ was overexpressed in BL21 (DE3) *E. coli* (TransGenBiotech, CD601-03) at 19°C for 16 h after induction by adding 50 µM of IPTG. Cells were harvested by centrifugation at 4000 rpm for 20 min at 4 °C and lysed with 50 mL of lysis buffer (50 mM Tris-HCl, 500 mM NaCl, pH 8.0, 10 mM imidazole, 4 mM β-mercaptoethanol, 1mM PMSF). After cell lysates were filtered with a 0.22 μm filter, protein was purified by Ni column (GE Healthcare, USA), eluted with elution buffer (50 mM Tris-HCl, 500 mM NaCl, pH 8.0, 250 mM imidazole and 4 mM β-mercaptoethanol), and proteins were further purified by Superdex 200 16/600 column (GE Healthcare) in a buffer containing 50 mM Tris-HCl pH 7.5, 300 mM NaCl and 2 mM DTT, and freshly frozen in liquid nitrogen and stored at −80 °C.

TDP-43^274-414^: TDP-43^274-414^ was expressed in BL21(DE3) *E. coli* (TransGenBiotech, CD601-03) into inclusion body. Cells were harvested and were lysed in a denatured lysis buffer (50 mM Tris-HCl, pH 8.0, and 6 M guanidine hydrochloride) at room temperature. Cell lysate was sonicated, followed by centrifugation at 14,000 rpm for 1 h at 4°C. Protein was purified from the supernatant by using a Ni column with the elution buffer containing 50 mM Tris-HCl at pH 8.0, 6 M guanidine hydrochloride and 50 mM imidazole. After further purification by HPLC (Agilent) with the elution buffer containing 35% (v/v) acetonitrile, the purified TDP-43^274-414^ was freeze-dried by FreeZone Lyophilizers (Thermo Fisher) and stored at −20°C.

hnRNP A1: hnRNP A1 was expressed in *E. coli* BL21 (DE3) pLysS after adding 0.4 mM IPTG at 25°C overnight. Cells were harvested and lysed in a lysis buffer (50 mM Tris-HCl at pH 7.5, 2 mM DTT, 1 mM PMSF and 5% glycerin) at 4 °C. The supernatant was loaded onto a 5 ml SP column by using a ÄKTA Purifier (GE Healthcare, USA). The proteins were eluted with a gradient mixing of buffer A (50 mM Tris-HCl pH 7.5, 2 mM DTT and 5% glycerin) and buffer B (buffer A with 1 M NaCl). Then, hnRNPA1 protein were further purified by Superdex 75 16/600 column (GE Healthcare) in a buffer containing 50 mM Tris-HCl pH 7.5, 500 mM NaCl and 2 mM DTT. Fractions containing hnRNPA1 monomers were collected and concentrated for following studies.

All the purified proteins were confirmed by Coomassie brilliant blue staining and Western blotting.

### *In vitro* PARylation assay

In vitro PARylation assay was performed according to the protocol adapted from (Slade et al., 2011). Briefly, substrate proteins were incubated with PARP1 (Sino Biological) in the presence of NAD in reaction buffer (50 mM Tris-Hcl, pH 7.4, 2 mM MgCl_2_) with or without 2.5 µg of ssDNA (Sigma) at 37 °C for 30 min. The reactions were stopped by adding 20 µM Olaparib and the products were examined by SDS-PAGE and Western blotting.

### *In vitro* phase separation

For the *in vitro* demixing experiments, purified hnRNP A1, PAR polymers (Trevigen, 4336-100-01) and NaCl at indicated concentrations were mixed into a phase separation buffer (50 mM Tris-HCl, pH 7.5, 10% (w/v) PEG (Sigma) and 2 mM DTT) and incubated for 3 min at room temperature. For co-LLPS, hnRNP A1 protein were incubated with TDP-43 in a buffer containing 50 mM Tris-HCl, pH 7.5, 100 mM NaCl and 2 mM DTT. 5 µL of each samples was pipetted onto a coverslip and imaged using a Leica microscope with differential interference contrast (DIC).

### Fluorophore-labeled of hnRNP A1 and TDP-43

Purified TDP-43^1-274^ and hnRNP A1 proteins in storage buffer were desalted in a reaction buffer (50 mM Tris-HCl, pH 7.5, 500 mM NaCl and 4 mM Tris (2-Carboxyethyl) Phosphine (TCEP) (Invitrogen, T2556) using a desalting column (GE Healthcare, USA) to remove DTT. The proteins were then incubated with 5-fold AlexaFluor-555 C2-malemide (invitrogen, A20346) for TDP-43^1-274^ or AlexaFluor-647 C2-malemide (invitrogen, A20347) for hnRNPA1 at room temperature for 2 h to conjugate the malemide derivative dye to a thiol group of Cys on the target proteins. The labeled proteins were further purified by Superdex 200 10/300 columns (GE Healthcare, USA) in a buffer containing 50 mM Tris-HCl, pH 7.5, 500 mM NaCl. Unlabeled proteins were mixed with one percent of labeled proteins for subsequent *in vitro* phase separation and confocal imaging.

### Lentivirus production and infection

293T cells were co-transfected with psPAX2, pMD2.G and pCDH-CMV-MCS-EF1-Puro plasmids of Flag, TDP-43-Flag, WT and the mutant hnRNP A1-HA (K298A and PBM^mut^), using the PolyJet™ reagent. Cell culture medium was collected and filtered through a 0.45 µm syringe filter (Millipore, SLHV033RB) at 48 h after transfection. The lentivirus was then concentrated using Lenti-X™ Concentrator (Clontech, PT4421-2). The 20-fold concentrated medium containing the desired lentivirus was used to infect NSC-34 cells in the subsequent experiments.

### Cell viability assay

NSC-34 cells were seeded into 96-well plates (Corning) at the density of 1.28×10^4^ cells/well and cultured in 100 µL of cultre medium containing the lentivirus particles for the expression of the proteins of interest. 48 h later, the infection medium containing lentivirus was removed and replaced with fresh medium. 48-72 h post infection, cell viability was examined using the Cell Counting Kit-8 (CCK-8) (Dojindo) according to the manufacturer’s instructions. Briefly, 10 µL of CCK-8 solution was added to each well. After incubation at 37°C for 2.5 h, the absorbance at at 450 nm was measured with a Synergy2 microplate reader (BioTek Instruments).

### TUNEL staining

TUNEL staining assay was performed using the TMR red *in situ* Cell Death Detection Kit (Sigma-Aldrich) according to the manufacturer’s instruction. The transfected NSC-34 cells grown on coverslips in a 24-well plate were fixed with 4% paraformaldehyde for 15 min and permeabilized with PBST (1xPBS + 0.5% Triton X-100) for 10 min at room temperature, followed by incubation with 18 µl labeling solution plus 2 µl enzyme solution at 37 °C for 1 h. Cell were mounted on glass slides using Vectashield Antifade Mounting Medium with DAPI (Vector Laboratories) and imaged using the Leica TCS SP8 confocal microscopy system.

### *Drosophila* strains

The following strains were obtained from the Bloomington *Drosophila* Stock Center (BDSC): RNAi-*Parp* (#57265), *elav*GS (#43642), RNAi-*mCherry* (#35785, a control for *in vivo* RNAi knockdown). The transgenic fly strain of UAS-TDP-43 was generated by ΦC31 integrase-mediated, site-specific integration into the fly genome, which allowed uniform transgene expression across different lines. The attP landing site stock used in this study was UAS-phi2b2a;VK5 (75B1) and a transgenic pBID-UASC-Luciferase (UAS-*Luc*) fly strain generated using the same method and the same landing site (Cao et al., 2017) was used as a control in this study. All flies were raised on standard cornmeal media and maintained at 25 °C and 60% relative humidity.

### Climbing ability and Lifespan assays

For the climbing assays, 20 flies per vial, 5-8 vials per group were tested. All flies were transferred into an empty polystyrene vial and gently tapped down to the bottom before 15 min adapted time for flies. The number of flies climbing over the height of 3 cm within 10 seconds was recorded. The test was repeated three times for each vial and 6∼8 vials per group were tested.

For the lifespan experiments, 20 flies per vial, 7-9 vials per group were tested. Flies were transferred to fresh fly food every 3 days. The flies lost prior to natural death because of escape or accidental death were excluded from the final analysis. The median lifespan was calculated as the mean of the medians of each vial in a group, whereas the “50% survival” shown on the survival curves is derived from compilation of all vials of a group. For adult-onset, neuronal expression of the RNAi transgenes using the *elav*-GS driver (Osterwalder et al., 2001), flies were raised at 25 °C and 60% relative humidity on regular fly food supplemented with 80 µg/ml RU486 (TCI).

### Statistical analysis

Unless otherwise noted, statistical significance in this study is determined by unpaired, two-tailed Student’s *t*-test at \**p* < 0.05, \*\**p* < 0.01, and \*\*\**p* < 0.001. Error bars represent the standard error of the mean (SEM).

## ACKNOWLEDGEMENTS

We thank the BDSC for providing fly strains, X. Gui for assistance in protein pruification, B. Yang for help in *in vitro* PARylation assay, S. Qiu and S. Zhang for technical supports, and J. Yuan, A. Li and members of the Liu lab and the Fang lab for helpful discussion and critical reading of the manuscript. This study was supported by grants from the National Key R&D Program of China (No. 2016YFA0501902) and the National Natural Science Foundation of China (NSFC, No. 31471017 and No. 81671254) to Y.F., the NSFC grants (No. 31470748) to C.L., (No. 21778063 and No. 91753114) to H.J., (No. 31500665 and No. 31671428) to Y.Z., and funding from the Shanghai Science and Technology Committee (No. 18ZR1448300) to A.D.

## AUTHOR CONTRIBUTIONS

YD, HJ, CL and YF conceived the research; YD, AD, YZ, HJ, CL and YF designed the experiments; YD, AD, JG, LS, GD, CW, ZM, LS, BQ, and KT performed the experiments; YD, AD, GD, CW, ZM, XD and HJ contributed important new reagents; YD, AD, JG, GD, CW, ZM, LS, KZ, and YZ analyzed the data; YD, AD, JG, LS, CL and YF interpreted and discussed the results; YD, AD, JG, GD and YF prepared the figures; and YD, AD, GD, CL and YF wrote the paper. All authors read and approved the final manuscript.

## SUPPORTING INFORMATION

Supporting information includes 7 figures.

